# Field evidence for manipulation of mosquito host selection by the human malaria parasite, *Plasmodium falciparum*

**DOI:** 10.1101/207183

**Authors:** Amélie Vantaux, Franck Yao, Domonbabele FdS Hien, Edwige Guissou, Bienvenue K. Yameogo, Louis-Clément Gouagna, Didier Fontenille, François Renaud, Frédéric Simard, Carlo Constantini, Fréderic Thomas, Karine Mouline, Benjamin Roche, Anna Cohuet, Kounbobr R Dabiré, Thierry Lefèvre

## Abstract

Whether the malaria parasite *Plasmodium falciparum* can manipulate mosquito host choice in ways that enhance parasite transmission toward humans is unknown. We assessed the influence of *P. falciparum* on the blood-feeding behaviour of three of its major vectors (*Anopheles coluzzii, An. gambiae* and *An. arabiensis*) in Burkina Faso. Host preference assays using odour-baited traps revealed no effect of infection on mosquito long-range anthropophily. However, the identification of the blood meal origin of mosquitoes showed that females carrying sporozoites, the mature transmissible stage of the parasite, displayed a 24% increase in anthropophagy compared to both females harbouring oocysts, the parasite immature stage, and uninfected individuals. Using a mathematical model, we further showed that this increased anthropophagy in infectious females resulted in a > 250% increase in parasite transmission potential, everything else being equal. This important epidemiological consequence highlights the importance of vector control tools targeting infectious females.

## Introduction

There is mounting evidence that malaria parasites affect phenotypic traits of their vectors and hosts in ways that increase contacts between them, hence favouring parasite transmission (Hurd 2003, Koella 2005, Lefèvre and Thomas 2008). In addition to increased vertebrate attractiveness to mosquito vectors (Lacroix *et al.* 2005, Cornet *et al.* 2013, Batista *et al.* 2014, De Moraes *et al.* 2014, Busula *et al.* 2017, Emami *et al.* 2017), another frequently reported parasite-induced change is the alteration of vector motivation and avidity to feed (Cator *et al.* 2012, Stanczyk *et al.* 2017). Mosquitoes infected with *Plasmodium* sporozoites (the mosquito to human transmission stage) can indeed display increased (i) responses to host odours (Rossignol *et al.* 1986, Cator *et al.* 2013), (ii) landing and biting activity (Rossignol *et al.* 1984, Rossignol *et al.* 1986, Wekesa *et al.* 1992, Anderson *et al.* 1999, Koella *et al.* 2002, Smallegange *et al.* 2013), (iii) number of feeds (Koella *et al.* 1998) and (iv) blood volume intake (Koella and Packer 1996, Koella *et al.* 1998, Koella *et al.* 2002). In contrast, mosquitoes infected with oocysts (the immature non-transmissible stage of the parasite), are less likely to attempt to feed (Anderson *et al.* 1999, Koella *et al.* 2002, Cator *et al.* 2013). Since biting is risky (e.g., host defensive behaviours can kill the vector and its parasite), reduced feeding attempts would be beneficial to the parasite during the non-transmissible stage as this would reduce mortality before the parasite reaches maturity and is ready to be transmitted (Schwartz and Koella 2001).

These “stage-dependent” behavioural alterations likely increase parasite transmission (Dobson 1988, Cator *et al.* 2014), provided that mosquito feeds are taken on a suitable vertebrate host species for the parasite. While malaria vectors can usually feed on a range of different vertebrate species (Takken and Verhulst 2013), the malaria parasites they transmit are often highly host-specific, infecting only one or a few vertebrate species (Perkins 2014). For example *P. falciparum*, which causes the most severe form of human malaria, displays an extreme form of specificity and can develop and reproduce in hominids only (predominantly in humans and to a lesser extent in chimpanzees, bonobos, and gorillas) (Prugnolle *et al.* 2011, Rayner *et al.* 2011, Ngoubangoye *et al.* 2016), such that any mosquito bite on another vertebrate species would be a dead-end for the parasite. In contrast, the vectors of *P. falciparum* can feed on a wide range of vertebrate host species in the wild depending on the geographic area and the relative abundance of humans and other vertebrates (Costantini *et al.* 1999, Takken and Verhulst 2013). Accordingly, *P. falciparum* could modify its vector choice in ways that enhance transmission toward humans and/or reduce mosquito attraction to other unsuitable host species (i.e. specific manipulation). A previous study testing this hypothesis found no effect of *P. falciparum* infection on host preference of three major vector species, *An. coluzzii*, *An. gambiae,* and *An. arabiensis* (Nguyen *et al.* 2017). However, this study examined the odour-mediated mosquito host preference in laboratory conditions using a dual-port olfactometer, not the final realised host choice which is of primary importance for parasite transmission.

Here, we assessed the influence of *P. falciparum* on *An. coluzzii, An. gambiae* and *An. arabiensis* blood-feeding behaviour in three villages in Burkina Faso. First, odour-baited traps, set side by side in a choice arrangement and releasing either human or calf odours were used to determine odour-mediated mosquito host preference (Experiment 1). Second, indoor-resting blood-fed mosquito females were collected and the origin of their blood meal was identified to determine mosquito host selection (Experiment 2). Third, we quantified the epidemiological consequences of variation in the patterns of host selection using a compartmental model for *Plasmodium* transmission between humans and mosquitoes.

## Material and methods

### Collection sites

The study was conducted in three villages in South-Western Burkina Faso: Soumousso (11°23’14”N, 4°24’42”W), Klesso (10°56’40.5”N, 3°59’09.9”W) and Samendeni (11°27’14.3”N, 4°27’37.6”W) (Figure supplement S1). The three villages are located in an area characterized by wooded savannah, where *Anopheles* females only have access to temporary, rain-filled puddles and quarries that permit larval development during the rainy season from June to November. The dry season extends from December to May. In these rural villages, domestic animals (including cattle, goats, sheep, pigs, chickens, donkeys, dogs) are usually kept in compounds in open conditions but a few households use separate roofed shelters for sheep, goats, pigs and chickens. Most houses are mud-walled with roofs of iron sheets or thatch, but a few houses are made of bricks.

### Experiment 1: Mosquito host preference

Two odour-baited entry traps (OBETs as in Costantini *et al.* 1996, Costantini *et al.* 1998, Lefèvre *et al.* 2009) and two odour-baited double net traps (BNTs as in Tangena *et al.* 2015) baited with calf and human odours were used to assess the host preference of field populations of mosquitoes in Samandeni and Klesso villages (Figure 1).The two OBETs were connected to a tent (Lxlxh: 250×150×150 cm) by air vent hoses (Scanpart^®^, DxL=10*300cm; Figure 1a). The odours of the two hosts were drawn by a 12-V fan from the tents and into the OBETs by the air vent hoses, coming out of the traps at a speed of 15cm/s (±2cm/s), as measured with a Testo 425-Compact Thermal Anemometer (Testo, Forbach, France) equipped with a hot wire probe [range: 0 to + 20m/s, accuracy: ± (0.03 m/s + 5% of mv)]. Host-seeking mosquitoes responding to the host cues flew up the odour-laden streams and entered one of the two traps. The two odour-baited double net traps (BNTs) consisted of an untreated bed net (Lxlxh: 300×250×185 cm) from which each corner was raised 20 cm above ground and a smaller untreated bed net (Lxlxh: 190×120×150 cm) protecting the human volunteer in the human baited trap (Figure 1b).

**Figure 1.**
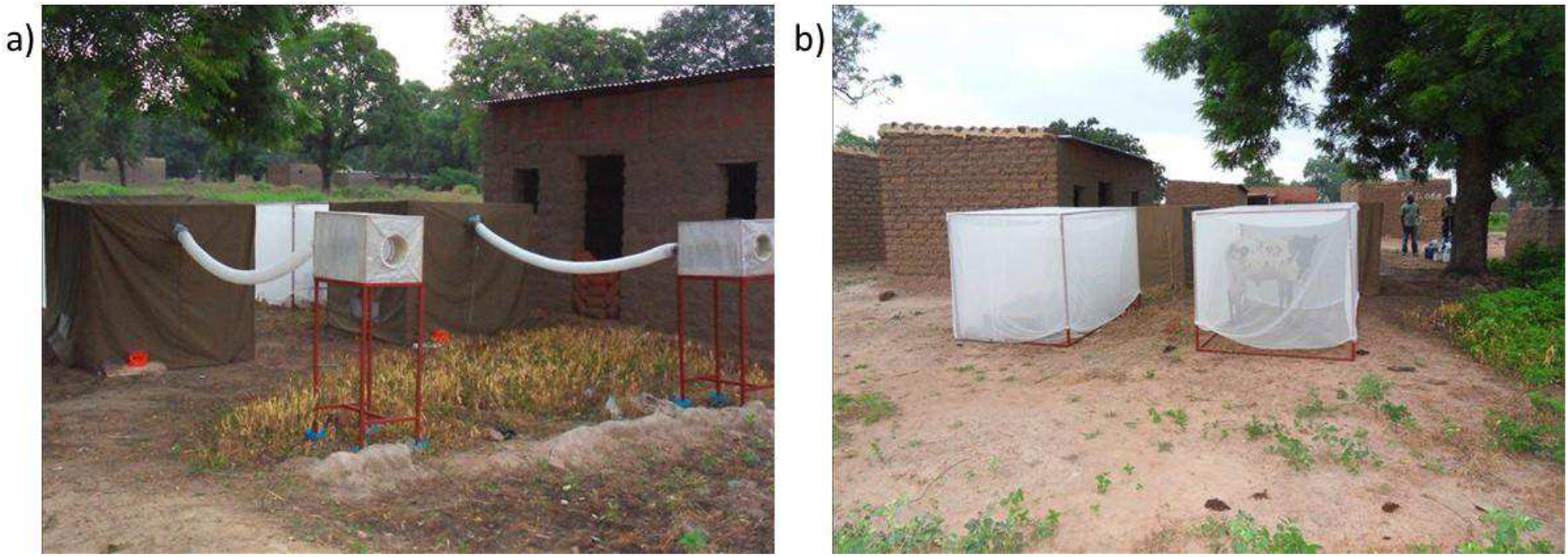
Traps baited with calf and human odours used to assess the host preference of field populations of mosquitoes in Samandeni and Klesso villages. **a)** Two odour-baited entry traps (OBETs) were connected to a tent by air vent hoses. **b)** Two odour-baited double net traps (BNTs).

In both OBETs and BNTs, the human volunteers rested on a metal-framed bed (Lxl: 190×80 cm) and were protected from mosquito bites. OBETs and BNTs were operated from 19:00 to 05:30 hours, for 3 nights in June 2013, and 13 nights in September 2013 in Samendeni. The BDNTs only were set-up for 6 nights in September in Klesso. Different combinations of live calves and humans were used as odour sources on each testing day to obviate any individual effect. Calves of about similar size and weight as human volunteers were used to equalize the quantity of emitted odours. Trapped mosquitoes were retrieved in the morning using mouth aspirators. They were kept in a 20×20×20 cm cage with a humid towel on top and brought back to the laboratory for further processing (see below).

### Experiment 2: Mosquito blood-feeding pattern

Indoor resting blood-fed mosquitoes were collected between 7 am and 9 am by insecticide spray catches as in Lefèvre et al. (2009) to determine the origin of their blood-meal. Briefly, white sheets were spread over the floor surface and the furniture inside houses. The houses were then sprayed with an insecticide (Kaltox^®^: allethrin 0.27%, tetramethrin 0.20 %, permethrin 0.17%, propoxur 0.68%) to knock down the mosquitoes. Fifteen minutes after spraying, blood-fed *An. gambiae s.l.* mosquitoes were collected from the white sheet using forceps and placed on moist filter paper inside labeled petri dishes.

In Samandeni and Klesso, mosquito collections were carried out in the rainy season only (4 days in June 2013, and 13 days in September 2013 in Samendeni, and 6 days in September 2015 in Klesso), whereas in Soumousso they were conducted in both the rainy and the dry season (26 days between January and November 2009). In Soumousso, human dwellings (from 10 neighbourhoods) only were sampled whereas animal sheds and unoccupied houses were also sampled in Samandeni and Klesso. A total of 27 human dwellings, 7 unoccupied houses and 20 animal sheds were sampled in Samendeni. A total of 7 human dwellings, 7 unoccupied houses and 9 animal sheds were sampled in Klesso. All mosquitoes were kept in a Petri dish with a humid paper towel to facilitate later dissection and brought back to the laboratory for further processing (see below).

### Laboratory processing of samples

A total of 3447 blood-fed *Anopheles gambiae s.l*. collected indoors (Experiment 2) and 674 females collected in the choice traps (Experiment 1) were processed. In addition, a subset of 276 females collected indoors was used to determine parity (parous versus nulliparous) based on the condition of ovarian tracheoles in order to control for age. Similarly, a subset of 418 individuals was used to determine different species within the *Anopheles gambiae sensu stricto* complex (i.e. distinguishing *Anopheles arabiensis*, *Anopheles coluzzii* and *Anopheles gambiae*) using routine PCR-RFLP based on segregating SNP polymorphisms in the X-linked ribosomal DNA InterGenic Spacer region as described in Santolamazza *et al*. (2008).

*Anopheles gambiae sl.* females were dissected in a drop of phosphate buffered saline (PBS) (pH 7.2). Blood-fed midguts were gently squeezed under a stereomicroscope (magnification 35x, Leica EZ4D, Wetzlar, Deutschland) to get the blood out, which was mixed with PBS, absorbed on a filter paper, and then kept at −20°C until identification by an enzyme-linked-immunosorbent assay (ELISA) for Soumousso and Samendeni samples (Beier *et al.* 1988) and by multiplex PCR for Klesso samples (Kent and Norris 2005). Each blood meal was discriminated between human, cattle, goat/sheep, chicken, dog, pig, and horse/donkey origins. ELISA-based determination of mosquito blood meal origin was performed using anti-human IgG-, anti-bovine IgG-, anti-pig IgG, anti-chicken IgG-, anti-goat IgG-, anti-sheep IgG-, anti-dog IgG-, and anti-horse IgG-peroxidase conjugates (A8794, A5295, A5670, A9046, A5420, A3415, A6792, A6917, Sigma-Aldrich). PCR-based determination of the mosquito blood meal origin targeting the vertebrate host cytochrome B was performed as described by Kent and Norris (2005), with the following modifications: *(i)* Three additional primers were designed from available Genbank sequences to target the following potential hosts: chicken470F (Genbank accession number: AB044986.1), sheep695F (KY662385.1), donkey574F (FJ428520.1); *(ii)* for each individual, two multiplex reactions were performed to avoid cross-reactions between primers and to optimize the determination. In the multiplex reaction #1, UNREV1025, Chicken470F, Sheep695F, Goat894F and Donkey574F primers were used at an amplification temperature of 49.2 °C. In the multiplex reaction #2, UNREV1025, Dog368F, Human741F, Cow121F and Pig573F primers were used at an amplification temperature of 58°C. Blood meal origin diagnostic was based on the PCR products expected sizes as follow: donkey (460bp), sheep (340bp), chicken (290bp), goat (150bp), dog (680bp), cow (561bp), pig (453bp), human (334bp).

The extracted midguts were then stained with 1% Mercurochrome^®^ solution to detect with a microscope (magnification 400x, Leica ICC50, Wetzlar, Deutschland) the presence and number of *Plasmodium* spp. oocysts. PCR on a subset of oocyst-infected individuals (20 midguts of a total of 118 oocyst-infected individuals) confirmed that these oocysts all belonged to *P. falciparum*. The head and thorax of individual mosquitoes were stored at −20°C in 1.5 mL Eppendorf tubes. Sporozoite infection with *P. falciparum* was determined by ELISA using peroxidase-conjugated *Plasmodium falciparum* circumsporozoite protein monoclonal antibody for the Soumousso samples (Wirtz *et al.* 1987) and by qPCR for the samples from Samendeni and Klesso (Boissière *et al.* 2013). The quantification of *P. falciparum* sporozoites in salivary glands was determined by qPCR using 7500 Fast Real time PCR System (Applied Biosystems, Foster City CA, USA). The mosquito heads and thoraxes were crushed individually and DNA extracted as previously described (Morlais *et al.* 2004). For sporozoite quantification, we targeted the fragment of subunit 1 of the mitochondrial cytochrome c oxidase gene (cox 1) using the forward and reverse primer sequences, qPCR-PfF 5’-TTACATCAGGAATGTTATTGC-3’ and qPCR-PfR 5’-ATATTGGATCTCCTGCAAAT-3, respectively. The reaction was conducted in a 10μL final volume containing: 1μL of DNA template, 1x HOT Pol EvaGreen qPCR Mix Plus ROX, and 600nM of each primer. Amplification was started by an initial activation step at 95°C for 15min and 40 cycles of denaturation at 95°C for 15s and annealing / extension at 58°C for 30s. Detection was conducted during the last step (Boissière *et al.* 2013). Quantification was based on a standard curve built from four serial dilutions (12%) of an asexual parasite culture. We made dilutions ranging from 60 to 60,000 genome/μl of DNAs from a standard culture. The first dilution (10^-1^) was used as a positive control. The standard curve (y=-3.384X +35.874) was obtained by linear regression analysis of Ct values (Cycle threshold) *versus* log10 genome copy number of parasite culture.

This protocol allowed us to gather the following information for each collected individual mosquito: immature *Plasmodium* infection status (presence of oocysts in the midgut); mature *P. falciparum* infection status (presence of sporozoites in salivary glands); source of blood meal or trap (calf/human) chosen; shelter type (human dwellings, unoccupied houses, animal sheds).

### Statistical analyses

#### Experiment 1: Mosquito host preference

The anthropophily index (AI) was expressed as the number of *Anopheles gambiae s.l.* caught in the human-baited trap over the total number of mosquitoes caught in both human- and calf-baited traps. We tested the effect of infection status (uninfected, infected with the oocyst immature stages and infected with the sporozoite transmissible stages), collection method (OBET *vs*. BNT), and their interaction on AI using a General Linear Model (GLM) with a binomial error structure.

#### Experiment 2: Mosquito blood-feeding pattern

The human blood index (HBI) was expressed as the number of *Anopheles gambiae s.l.* fed on humans including mixed human-animal blood meals over the total number of blood-fed *Anopheles gambiae s.l.*. We tested the effect of *Plasmodium* infection status (uninfected, oocyst-infected, sporozoite-infected individuals - 25 individuals with both oocysts and sporozoites were included in the sporozoite infected group and excluding these individuals from the analysis yielded similar results), village (Soumousso, Samendeni, Klesso), shelter type (human dwelling, unoccupied house, animal shed) and relevant two-way interactions (infection status by shelter type and infection status by village) on HBI using a GLM with a binomial error structure. The effect of species (*Anopheles gambiae*, *An. coluzzii* and *An. arabiensis*), infection status, shelter type, and their interactions on HBI was assessed using the subset of females identified to the molecular level using a GLM with a binomial error structure. The effect of parity (nulliparous *vs*. parous) on HBI was assessed on a subset of females using a GLM with a binomial error structure.

We also verified for both AI and HBI whether choice significantly differed from a random distribution between humans and animals or whether mosquitoes displayed a statistically significant attraction to one type of blood meal or trap.

For model selection, we used the stepwise removal of terms, followed by likelihood ratio tests (LRT). Term removals that significantly reduced explanatory power (P<0.05) were retained in the minimal adequate model (Crawley 2007). All analyses were performed in R v.3.0.3.

### Mathematical model

In order to explore the epidemiological consequences of variation in HBI, we built a compartmental model for *Plasmodium* transmission between humans and mosquitoes (Keeling and Rohani 2008):

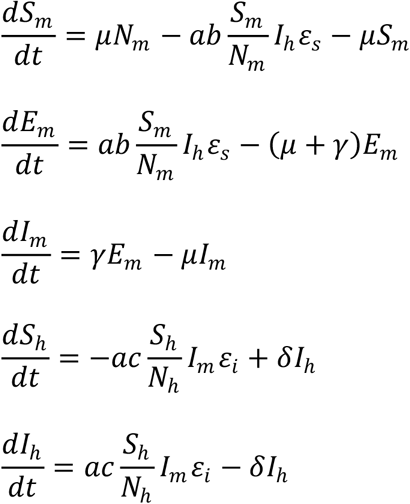

Susceptible mosquitoes (*S_m_*) are born at rate *μ* and become exposed (*E_m_*) according to their biting rate (*a*), their probability to get infected (*b*) and the HBI of susceptible mosquitoes (εs). Then, exposed mosquitoes become infectious (*I_m_*) according to their extrinsic incubation period (*γ*). Mosquito population die at rate (*μ*). Nm is the number of mosquitoes. Susceptible humans (*S_h_*) get infected according to mosquito biting rate, the probability to develop infection (c) and the HBI of infectious mosquitoes (ε_i_). Nh is the number of humans. Then, infectious humans remain infectious (*I_h_*) during a period equals to *1/δ* on average. See parameter values in table supplement S1 (Roux *et al.* 2015, Vantaux *et al.* 2016). In our simulation we based the HBI of exposed mosquitoes (*εs*) on the confidence intervals of oocyst-infected mosquitoes that were experimentally measured in this study. Then we explored the impact of the HBI of infectious mosquitoes (*εi*, during the sporozoite stage) on the Entomological Inoculation Rate (EIR), representing the number of infectious bites received by a human during one year (Smith and Ellis McKenzie 2004), as defined by:

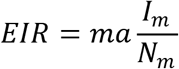

where *m* is the ratio between mosquitoes and humans, and other parameters are as above. We kept an identical human population size of 100 individuals and only varied mosquito densities to assume different ratio values (m) between mosquitoes and humans (low: m=1, medium: m=10 and high: m=100) in order to explore the impact of different HBIs on the EIR in relation to mosquito densities. Then, the mathematical model was simulated for one season in order to estimate the proportion of infectious mosquitoes.

### Ethics

Ethical approval was obtained from the Centre Muraz Institutional Ethics Committee under agreement no. 0003-2009/CE-CM and A0003-2012/CE-CM.

## Results

### Experiment 1: Mosquito host preference

To assess the inherent mosquito host preference of field populations of mosquitoes, we used two odour-baited entry traps (OBETs) and two odour-baited double net traps (BNTs) releasing either calf or human odours. The anthropophily index (AI) was expressed as the number of *Anopheles gambiae s.l.* caught in the human-baited trap over the total number of mosquitoes caught in both human- and calf-baited traps. The infection status was successfully determined in 584 out of the 674 mosquitoes (86.6%) collected in the OBETs (383 individuals) and BNTs (201 individuals). Uninfected, oocyst-infected and sporozoite-infected females displayed similar host preferences (X^2^_2_ = 3.6, P = 0.17, Figure supplement S2, AI uninfected females: 63.3 ± 4%, N=531, OR=0.58, 95% CI = 0.53-0.63, P <0.0001; AI oocyst-infected females: 55.2 ± 18 %, N=29, OR=0.81, 95% CI = 0.56-1.18, P=0.58; AI sporozoite-infected females: 45.8 ± 20 %; N=24, OR=1.18, 95% CI = 0.78-1.78, P=0.7). There was no effect of collection method on AI (OBETs: 64 ± 5%, BNTs: 59 ± 7%; X^2^_1_ = 1.5, P = 0.21), indicating that both methods are comparable to assess mosquito host preference. There was no interaction between mosquito infection and collection method (X^2^_2_ = 0.26, P = 0.9; Figure supplement S2).

### Experiment 2: Mosquito blood-feeding pattern

To assess the realized host selection of *Anopheles gambiae s.l.*, the blood meal origins of indoor-resting females were identified. The human blood index (HBI) was expressed as the number of females fed on humans (including mixed human-animal blood meals) over the total number of blood-fed females. Of the 3447 blood-fed *Anopheles gambiae s.l*. collected indoors, the blood meal origin was successfully identified in 2627 samples (76%). Among these 2627 samples, infection status was successfully determined in 2328 mosquitoes (88.6%). The following analyses are restricted to these 2328 females. HBI was significantly affected by mosquito infection status (X^2^_2_ = 13.007, P = 0.0015; Figure 2) with a 24% increase in HBI in sporozoite-infected females compared to both their oocyst-infected and uninfected counterparts (sporozoite-infected: 77 ± 5.7%; N=209, deviation from random feeding: OR=0.3, 95% CI = 0.25-035, P <0.0001; oocyst-infected females: 63.6 ± 5.7%, N=118, OR=0.57, 95% CI = 0.47-0.69, P =0.004; uninfected females: 61.1 ± 2.1%; N=2001, OR=0.64, 95% CI = 0.61-0.66, P <0.0001). However, because sample size in the uninfected group (N=2001) was higher than that of both sporozoite-infected (N= 209) and oocyst-infected groups (N=118), we ran a second set of analyses using a subset of 150 randomly selected uninfected individuals. This approach normalizes statistical power to test for statistically significant differences in HBI across heterogeneous sample sets. The randomisation was repeated 100 times and the analysis confirmed a significantly higher anthropophagy in sporozoite-infected individuals compared to both oocyst-infected individuals and uninfected individuals in 100% of these randomisations (mean (X^2^_2_) = 12.7, CI (X^2^_2_) = (7.54-21.59), mean (P) = 0.0043, CI(P) = (0.00002-0.023); Tukey post-hoc tests: sporozoite-infected *vs*. oocyst-infected individuals, this pair-wise comparison was significantly different in 100 % of the randomisations: mean(P) = 0.02577, CI(P) = (0.02559-0.02591); sporozoite-infected *vs*. uninfected individuals, this pair-wise comparison was significantly different in 90% of the randomisations: mean (P) = 0.023, CI(P) = (5e-07 - 3e-01); oocyst-infected *vs*. uninfected individuals, this pair-wise comparison was significantly different in 0 % of the randomisations: mean (P) = 0.78, CI(P) = (0.07-0.99)).

**Figure 2.**
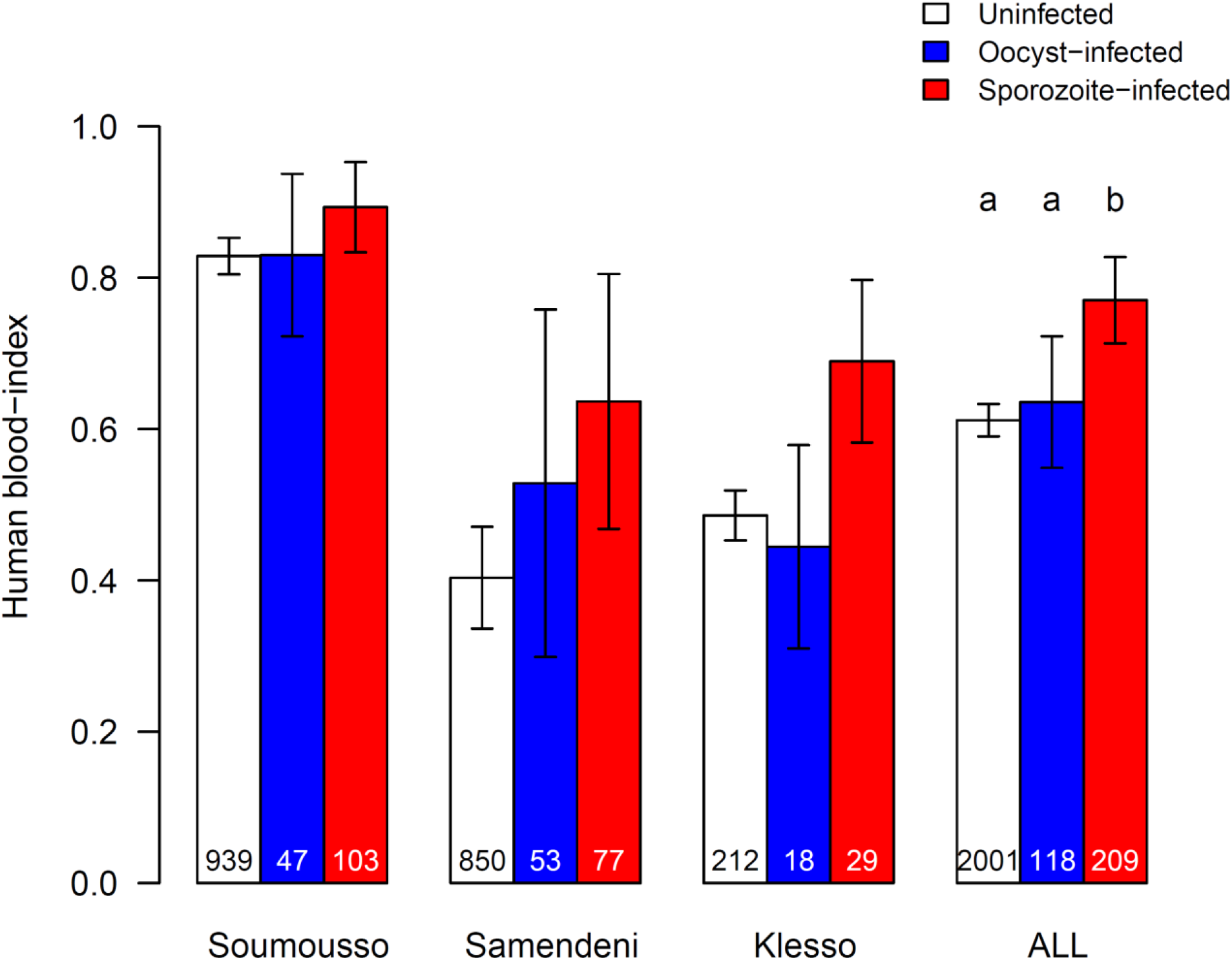
Effect of infection status on the human-blood index of *Anopheles gambiae s. l*. females expressed as the number of females fed on humans out of the total number of blood-fed females for the three sampled villages. Data show proportion ± 95% confidence intervals. Numbers in bars indicate the total numbers of mosquitoes. Different letters indicate differences between infection status (Chi-square post-hoc tests: sporozoite-infected *vs*. oocyst-infected females X^2^_1_=6.1, P=0.013; sporozoite-infected *vs*. uninfected females X^2^_1_=19.4, P<0.0001; oocyst-infected *vs*. uninfected females X^2^_1_=0.18, P= 0.67).

The HBI of sporozoite-infected mosquitoes was higher than that of oocyst-infected and uninfected females regardless of the village considered (infection status: village interaction: X^2^_4_ = 2.3, P = 0.68, Figure 2) or the shelter type in which mosquito females were collected (infection status: shelter type interaction: X^2^_4_ = 0.7, P = 0.95, Figure supplement S3).

HBI was also significantly influenced by shelter type (X^2^_2_ = 145.92, P < 0.0001). Females collected in animal sheds were significantly less likely to have fed on human hosts (22.3 ± 4%) than females collected in unoccupied houses (40.9 ± 6.8%; Chi-square post-hoc test: X^2^_1_ = 21.6, P < 0.0001) or in human dwellings (74.5 ± 2%; Chi-square post-hoc test: X^2^_1_ = 385, P < 0.0001). Females collected in human dwellings were also significantly more likely to have fed on human hosts than females collected in unoccupied houses (Chi-square post-hoc test: X^2^_1_ = 96, P < 0.0001). HBI was significantly affected by the village (X^2^_2_ = 139.5, P < 0.0001). However, in Soumousso only human dwellings were sampled confounding the effect of village and shelter type in this case. Therefore, we carried out an analysis on the human dwellings only to compare HBIs in the three villages. Mosquitoes were significantly less anthropophagic in Samendeni (56.5± 4%), compared to Soumousso (83.5±2.2%; Chi-square test: X^2^_1_ =138.8, P < 0.0001) and Klesso (77.3±9 %; Chi - square test: X^2^_1_ = 12.7, P = 0.0004). HBIs in Soumousso and Klesso were not significantly different (83.5±2.2% *vs*. 77.3±9 % respectively; Chi-square test: X^2^_1_ = 1.8, P = 0.18).

A significant species variation in HBI was observed (X^2^_2_ = 10.2, P = 0.006; Figure 3) with *Anopheles arabiensis* being significantly less anthropophagic (22.2 ± 15%, N=27, OR=3.5, 95% CI = 2.2-5.56, P = 0.007) than *An. gambiae* (54.8 ± 7.1%; N=186, OR=0.82, 95% CI = 0.71-0.95, P = 0.19) and *An. coluzzii* (55.1 ± 6.8%; N=205, OR=0.81, 95% CI = 0.71-0.94, P=0.14). Although HBI varied among mosquito species, sporozoite-infected individuals displayed the highest anthropophagy regardless of the species considered (infection status: species interaction: X^2^_4_ = 4, P = 0.42; Figure 3 and supplementary material).

**Figure 3.**
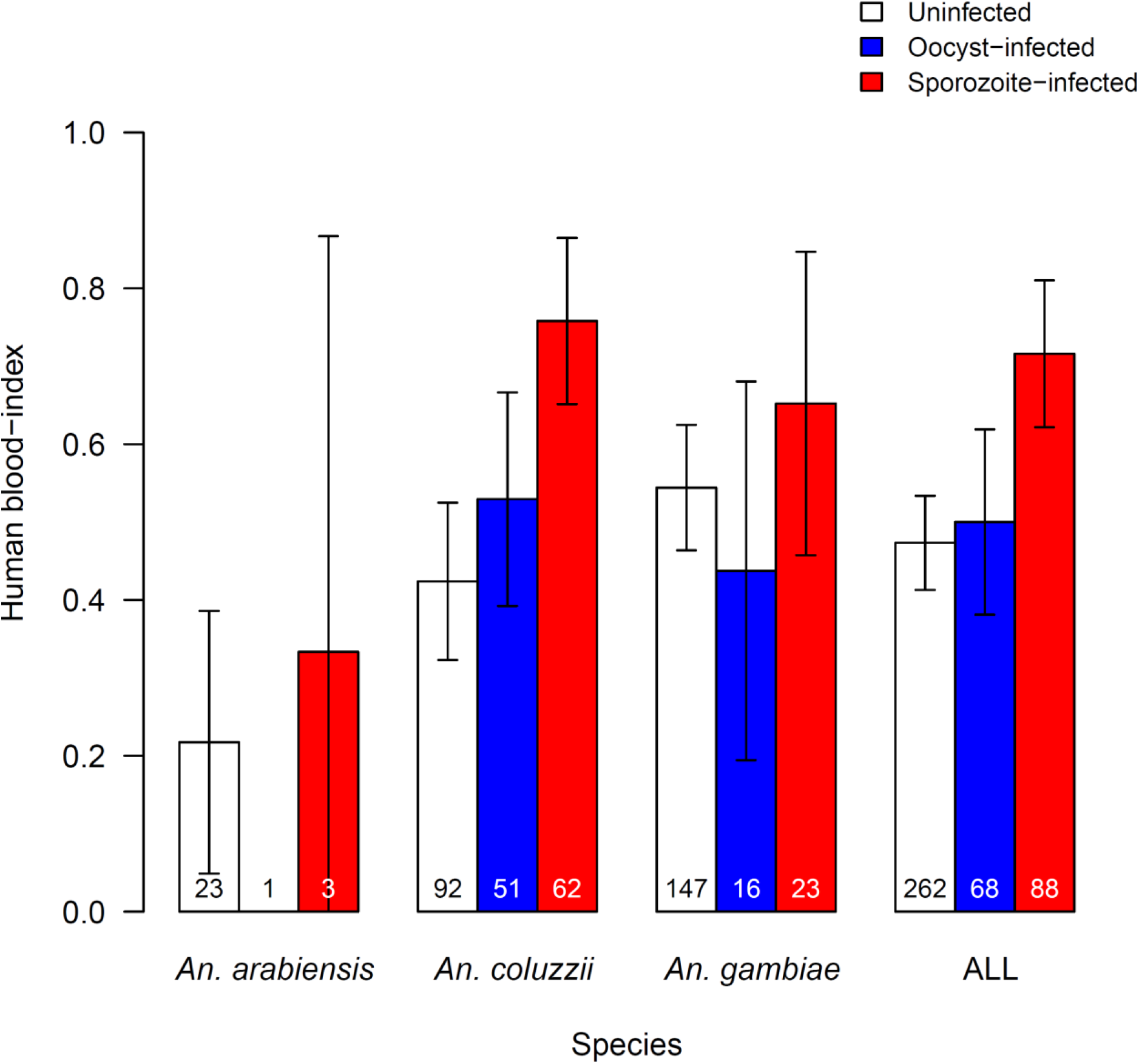
Effect of infection status and mosquito species on the human-blood index expressed as the proportion of females fed on humans or humans and animals out of the total of blood-fed females. Data show proportion ± 95% confidence intervals. Numbers in bars indicate the total numbers of mosquitoes.

Finally, HBI was not significantly affected by parity, a proxy used to estimate mosquito age (nulliparous females: 49.53 ± 9%, parous females: 45.6 ± 7.5%; X^2^_1_ = 0.4, P = 0.52).

### Epidemiological consequences

To investigate the epidemiological impact of a higher HBI in infectious females compared to oocyst-infected and uninfected females, we built a mathematical model based on the experimental values observed in this study. This model assessed the impact of different HBIs on the Entomological Inoculation Rate (EIR, number of infectious bites received by a person during one year) at different mosquito lifespans and densities. In order to consider the heterogeneity of HBI values on epidemiological consequences, the HBI of susceptible mosquitoes was based on the average value whereas the HBI of exposed mosquitoes were assumed to be uniformly distributed within the confidence intervals of the HBI of oocyst-infected mosquitoes that were experimentally measured in this study. Then, the impact of HBI variation in infectious (sporozoite-infected) mosquitoes on parasite transmission potential was explored fully (Figure 4). For an average mosquito lifespan of 15 days (Figure 4a), an HBI of 0.62 in infectious mosquitoes (similar to that of susceptible mosquitoes) resulted in an EIR of 4 at a low ratio of 1 (1 mosquito per human), while an HBI of 0.77 (as observed here in infectious mosquitoes) resulted in an EIR of 14. In other words, a 24% increase in HBI resulted in a 250% increase in EIR, everything else being equal. Transmission consequences were even larger when the human-to-mosquito ratios were higher (EIR = 5 *vs*. EIR = 19 with a ratio of 10 or 100, i.e. a 280% increase in EIR) but the size of the increase in EIR for sporozoite-infected mosquitoes declined with increasing mosquito longevity (Figure 4c, 4d, and supplementary material).

**Figure 4.**
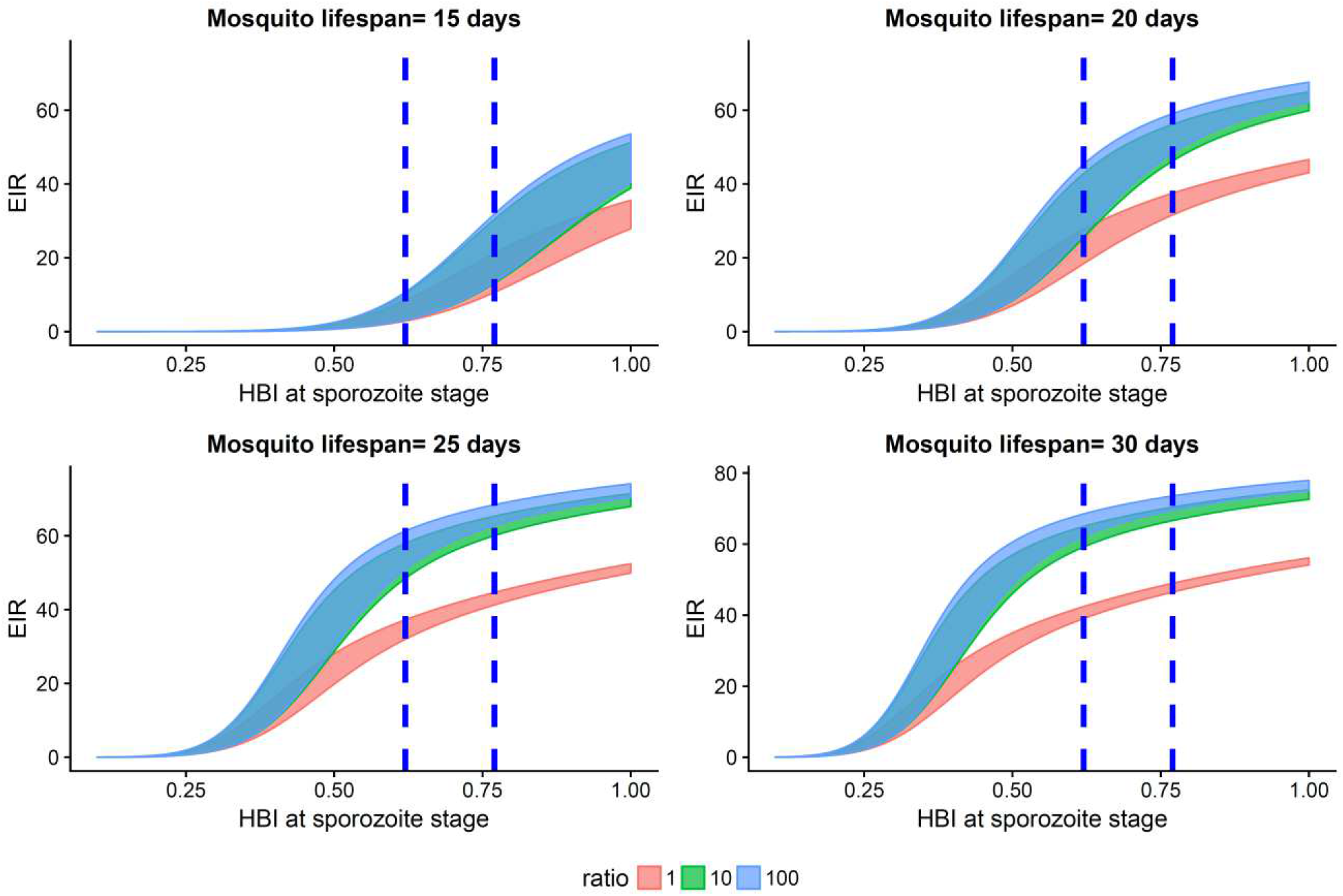
Expected epidemiological consequences of HBI variation for different values of mosquito lifespan and mosquito/human ratio. The X axis represents the range of values considered for the HBI of infectious (sporozoite-infected) mosquitoes and the Y axis is the Entomological Inoculation Rate (EIR, number of infectious bites received by a person over one year) when the HBI of exposed mosquitoes corresponds to the confidence intervals of the HBI of oocyst-infected mosquitoes that were experimentally measured in this study. The ribbons represent the possible EIR values for different HBI of sporozoite-infected mosquitoes according to the confidence interval of HBI in oocyst-infected mosquitoes (63.6% ± 5.7%) and for different values of the mosquito to human ratio. The dashed lines represents the average value measured for susceptible mosquitoes (0.62) and for sporozoite-infected mosquitoes (0.77). Ratio=adult mosquito/human densities.

## Discussion

The mosquito host preference assays (experiment 1 using OBETs and BNTs,) showed that infected mosquitoes displayed similar long-range attraction toward human odour as uninfected individuals regardless of parasite developmental stages (oocyst *vs*. sporozoite), confirming previous laboratory results (Nguyen *et al.* 2017). However, consistent with the hypothesis of specific manipulation, the patterns of mosquito host selection (experiment 2 based on identification of mosquito blood-meal sources) showed that sporozoite-infected *An. coluzzi*, *An. gambiae* and *An. Arabiensis* females were more likely to have fed on human than oocyst-infected and uninfected individuals. By distinguishing sporozoite and oocyst infection, we ruled out the potential confounding effect of a mere intrinsic mosquito characteristic. Infected mosquitoes may indeed exhibit increased anthropophagy not because of being infected but just because of an innate preference for humans, thus making these mosquito individuals infected. Here, individuals infected with sporozoites displayed different HBI than individuals infected with oocysts, thus ruling out this possibility. Because *Plasmodium falciparum* takes about 10 to 18 days to complete its development (depending on temperature, (Nikolaev 1935, Shapiro *et al.* 2017, Ohm *et al.* 2018) there is an increased likelihood of sporozoite infection as mosquitoes become older. This means that mosquito age could be a confounding factor of infection, with infected mosquitoes displaying increased HBI not because they harbour sporozoites but because they are older. Such an age effect could be mediated by specific physiological requirements in old mosquitoes or by a positive reinforcement (learning / memory) of feeding on humans. Our data does not support an age effect as we did not find a significant effect of parity (a proxy for age) on HBI (i.e. parous and nulliparous mosquito females displayed similar anthrophagy).

The precise mechanisms responsible for increased anthropophagy in sporozoite-infected mosquitoes is not yet clear, but at least three hypotheses can be proposed. First, malaria parasites might manipulate mosquito short-range behaviours only, whereas at longer range when mosquitoes rely mainly on CO_2_ and other volatile odours (Gillies 1980, Mboera and Takken 1997, Gibson and Torr 1999, Cardé and Gibson 2010), sporozoite-infected mosquitoes display similar preferences to uninfected and oocyst-infected individuals. At short range, mosquitoes rely on other cues including visual stimuli, moisture, heat and skin emanations (Gibson and Torr 1999, Cardé and Gibson 2010, Takken and Verhulst 2013). These stimuli can be host specific, and inform of host suitability for parasite development before the mosquito engages in selection and eventually in feeding. In addition to a possible preferential short-range attraction of sporozoite-infected mosquitoes toward host species suitable for parasite development, there could also be short-range repellence by unsuitable host species.

Second, the parasite may induce changes in the vector such as an alteration of microhabitat choice to spatially match the habitat of the suitable host. This could be achieved through parasite manipulation of mosquito endophagic/philic behaviours resulting in a higher degree of indoor-feeding and -resting of sporozoite-infected females. For example, infectious mosquitoes may exhibit an enhanced tendency to enter (or a decreased tendency to exit) house interstices regardless of emitted odours.

Third, the parasite may induce changes in the vector such as an alteration of time activity in order to temporally match the time of rest or activity of the suitable host. Mosquitoes exhibit circadian rhythms in many activities such as flight, host-seeking, swarming, egg-laying, etc. (Rund *et al.* 2016). There is mounting evidence that, following bed-net introduction, malaria vectors can display an increased tendency to feed outdoors (Russell *et al.* 2011) or bite earlier in the evening or later in the morning (Moiroux *et al.* 2012). Accordingly, *P. falciparum* could manipulate mosquito host-seeking rhythms in a way that increases bites on unprotected people. Testing this hypothesis would require sampling mosquitoes at distinct periods and comparing the proportion of uninfected, oocyst-infected and sporozoite-infected vectors among samples.

Sporozoite-induced change in mosquito host selection occurred in three major and related mosquito vectors, namely *An. coluzzii*, *An. gambiae* and *An. arabiensis*. This suggests that manipulation likely already occurred in the common ancestor of these three species and that the parasites might exploit a physiological pathway common to all three mosquito species to modify its vector host choice.

Transmission models generally assume that uninfected and infected vectors have similar preferences for human (Smith and Ellis McKenzie 2004, Smith *et al.* 2012). This study suggests that this assumption may not be valid and that these models possibly underestimate transmission intensity. Our modelling approach confirms that HBI increases in infectious mosquitoes can have a dramatic impact on disease transmission. In particular, if we consider mosquito lifespans relevant to natural settings (i.e. 15 to 20 days; Gillies 1961, Gillies and Wilkes 1965, Saul *et al.* 1990, Charlwood *et al.* 1997, Killeen *et al.* 2000), the transmission potential was almost multiplied by 3 when the HBI increased from 0.62 to 0.77 *i.e*. the value observed for the infectious mosquitoes in this study. For many mosquito–*Plasmodium* associations including *An. gambiae s.l.-P. falciparum*, the duration of the parasite’s development within the mosquito is as long as the insect vector’s average lifespan (Gillies 1961, Gillies and Wilkes 1965, Saul *et al.* 1990, Charlwood *et al.* 1997, Killeen *et al.* 2000, WHO 2014). This means that most mosquitoes do not live long enough to transmit the disease, and hence that feeds taken by infectious mosquitoes on unsuitable host species would have disastrous consequences for parasite fitness. The model suggests that the benefits of specific manipulation should be particularly high in vectorial systems in which transmission opportunities are rare (short vector lifespan, relatively long parasite development period, and diverse blood sources).

In conclusion, our results suggest that the human malaria parasite *P. falciparum* evolved the ability to enhance transmission toward humans, the appropriate host species, by increasing mosquito anthropophagy (or decreasing zoophagy) with potentially profound public health consequences. Future laboratory and field studies will be essential to confirm these results and to better understand the epidemiological, ecological and evolutionary consequences of parasite manipulation of vector behaviours.

## Acknowledgements

We would like to thank all volunteers for participating in this study as well as the local authorities for their support. We are very grateful to the IRSS staff in Burkina Faso for technical assistance. We thank Priscille Barreaux for discussion and comments. We are grateful to Alison Duncan, Ricardo S. Ramiro, Olivier Restif and an anonymous reviewer for comments and corrections which greatly improved the manuscript. This preprint has been reviewed and recommended by Peer Community In Evolutionary Biology (https://dx.doi.org/10.24072/pci.evolbiol.100057).

## Data availability

Raw data are available on zenodo: https://doi.org/10.5281/zenodo.1296744. Statistical analyses are available as supplementary file.

## Competing interests

We have no competing interests.

## Funding

This study was supported by the ANR grant no. 11-PDOC-006-01 and the European Community’s Seventh Framework Program (FP7/2007–2013) under grant agreements no. 242095 and no.223736. BR is supported by the ANR project PANIC.

